# Exaggerated Cortical Representation of Speech in Older Listeners: Mutual Information Analysis

**DOI:** 10.1101/2019.12.18.881334

**Authors:** Peng Zan, Alessandro Presacco, Samira Anderson, Jonathan Z. Simon

## Abstract

Aging is associated with an exaggerated representation of the speech envelope in auditory cortex. The relationship between this age-related exaggerated response and a listener’s ability to understand speech in noise remains an open question. Here, information-theory-based analysis methods are applied to magnetoencephalography (MEG) recordings of human listeners, investigating their cortical responses to continuous speech, using the novel non-linear measure of phase-locked mutual information between the speech stimuli and cortical responses. The cortex of older listeners shows an exaggerated level of mutual information, compared to younger listeners, for both attended and unattended speakers. The mutual information peaks for several distinct latencies: early (∼50 ms), middle (∼100 ms) and late (∼200 ms). For the late component, the neural enhancement of attended over unattended speech is affected by stimulus SNR, but the direction of this dependency is reversed by aging. Critically, in older listeners and for the same late component, greater cortical exaggeration is correlated with decreased behavioral inhibitory control. This negative correlation also carries over to speech intelligibility in noise, where greater cortical exaggeration in older listeners is correlated with worse speech intelligibility scores. Finally, an age-related lateralization difference is also seen for the ∼100 ms latency peaks, where older listeners show a bilateral response compared to younger listeners’ right-lateralization. Thus, this information-theory-based analysis provides new, and less coarse-grained, results regarding age-related change in auditory cortical speech processing, and its correlation with cognitive measures, compared to related linear measures.

**New & Noteworthy:** Cortical representations of natural speech are investigated using a novel non-linear approach based on mutual information. Cortical responses, phase-locked to the speech envelope, show an exaggerated level of mutual information associated with aging, appearing at several distinct latencies (∼50, ∼100 and ∼200 ms). Critically, for older listeners only, the ∼200 ms latency response components are correlated with specific behavioral measures, including behavioral inhibition and speech comprehension.

## Introduction

Young normal hearing listeners are capable of separating attended speech from background distractions, but this capability degrades with aging. Behavioral studies have shown age-related temporal processing deficits in a variety of auditory tasks, including pitch discrimination (Fitzgibbons and Gordon-Salantt 1996), gap-in-noise detection (Fitzgibbons and Gordon-Salant 2001) and recognition of speech in noise (Frisina and Frisina 1997; Gordon-Salant et al. 2006; He et al. 2008). Neurophysiological studies show that although the young auditory brain robustly segregates speech from either a competing speaker (Ding and Simon 2012a) or spectrally matched noise (Ding and Simon 2013), temporal aspects of neural processing demonstrate age-related changes in response latency and strength, in both midbrain (Anderson et al. 2012; Burkard and Sims 2002; Clinard and Tremblay 2013) and cortical evoked responses (Herrmann et al. 2019; Lister et al. 2011; Presacco et al. 2016a, 2016b). In animal studies, age-related increases in both spontaneous and stimulus-driven firing rates have been reported in the auditory cortex (Engle and Recanzone 2013; Hughes et al. 2010; Juarez-Salinas et al. 2010; Ng and Recanzone 2018; Overton and Recanzone 2016). In aging rats, altered inhibition and functional impairments in the cortex can arise from regulated plasticity change, and may be reversible (de Villers-Sidani et al. 2010). However, it remains an open question how much such plasticity change occurs in the aging human brain, and the extent of its effects on speech processing.

The MEG studies of Presacco et al. (2016a, 2016b), using a stimulus reconstruction paradigm, found an exaggerated response to speech in noise for older listeners by demonstrating a higher speech envelope reconstruction accuracy in older listeners than younger. A later re-analysis of the same data (for speech without noise) found that a major source of the exaggerated response is from response components with ∼50 ms latency; contributions from later latencies could not be ruled out but were not significant (Brodbeck et al. 2018). Response components with ∼100 ms latency are natural candidates since they are strongly attention-dependent (Ding and Simon 2012a, 2013), and older listeners might exert more attention than younger listeners. Also, since multi-modal association (binding) of auditory and visual responses occurs at latencies beyond the 100 ms (Griffiths and Warren 2004), we might also expect further contributions from later responses, for older listeners. Based on these previous findings, we hypothesize that older listeners will exhibit a higher level of mutual information than younger listeners for response components of 50 ms, 100 ms and even later latencies. Additionally, Presacco et al. (2016b) demonstrated a negative correlation between speech envelope reconstruction accuracy and a behavioral inhibition score (a visual flanker task) in older listeners, but it remains unknown which response latencies underlie this association.

In terms of hemispheric lateralization of cortical representations of speech, the results of Cabeza (2002) support a general reduction of lateralization in older adults for cognitive processing, including memory, attention and inhibitory control, denoted HAROLD (hemispheric asymmetry reduction in older adults). Here we investigate whether there might exist an analogous age-related lateralization change in speech processing, again using mutual information.

Investigations of cortical coding of continuous speech often rely on linear methods (Ding and Simon 2012a; Presacco et al. 2016a, 2016b). Auditory cortex, however, is well known to employ non-linear processing (Sahani and Linden 2003), and therefore a non-linear analysis framework may provide more insight. Nonlinear approaches based on Shannon’s information theory (Shannon 1948) have been successfully applied in the auditory system to spiking neurons (Nelken and Chechik 2007) and EEG subcortical recordings (Zan et al. 2019). Information theoretic approaches have also been applied to MEG recordings from auditory cortex (Cogan and Poeppel 2011), to decode phase information in low-frequency responses to speech. Additionally, by analyzing the mutual information between auditory midbrain and cortical responses, it can be seen that older listeners display redundant information during a task involving categorical perception of speech syllables (Bidelman et al. 2014).

Here, to investigate the information encoded in cortical responses phase-locked to continuous speech, we develop the temporal mutual information function (TMIF) measure. It provides a novel non-linear measure of a general phase-locked response to speech, analogous to the linear temporal response function (TRF), or (linearly averaged) evoked responses to a brief sound. Like both, it also has response components with peaks at specific latencies, analogous to the TRF’s M50TRF and M100 TRF components, or the M50 and M100 response components of an evoked response. The main mutual information peaks of the TMIF are, by analogy, named the MI50, MI100 and MI200, and occur for early cortical latency (∼50 ms), middle cortical latency (∼100 ms), and late cortical latency (∼200 ms).

## Materials and methods

### Subjects

The dataset analyzed here was previously obtained and analyzed in earlier studies (Brodbeck et al. 2018; Presacco et al. 2016a, 2016b). 32 subjects participated in the experiment: 17 younger adults ages 18 to 27 (3 male) and 15 older adults ages 61 to 73 (5 male). All participants were recruited from the greater Washington D.C. area (Maryland, Virginia and Washington D.C.), with clinically normal hearing. Specifically, participants had normal hearing thresholds (≤ 25 dB hearing level) from 125 Hz to 4000 Hz, no history of neurological or middle ear disorders or surgery, and normal intelligent quotient scores [≥ 85 on the Wechsler Abbreviated Scale of Intelligence (Zhu and Garcia 1999)]. Written informed consent was obtained from each subject, and they were compensated for their time. The experimental protocol and all procedures were reviewed and approved by the Institutional Review Board of the University of Maryland.

### Behavioral tests

#### Flanker test

The ability to attend to a selected or goal-appropriate stimulus and to ignore other distracting stimuli is associated with inhibitory control (Neill et al. 1995), and this ability declines with aging (Diamond 2013). This ability may affect auditory suppression of a competing speaker while attending to another. To investigate broad aging effects on behavioral inhibition, including its relationship with complex auditory processing, a visual Flanker test (Ward et al. 2016) was given to all subjects. The Flanker test measured behavioral inhibition and attention control by displaying five arrows in a row and asking only for the direction of the middle arrow, i.e., the flanking arrows serve only as distractors. Both reaction time and accuracy are taken into account for scoring (Weintraub et al. 2013), and a *higher* Flanker score indicates better performance, i.e., more control of behavioral inhibition.

#### QuickSIN test

The Quick Speech-in-Noise test (QuickSIN) measures listeners’ ability to understand speech in noise (four-speaker babble), with subjects asked to recall words presented at six signal-to-noise ratio (SNR) levels (ranging from 0 dB to 25 dB SNR), with performance rated by the number of key words they correctly recalled (Killion et al. 2004). An SNR loss is calculated from the total number of key words correctly repeated. A *lower* QuickSIN SNR loss indicates better performance, i.e., superior ability to understand speech in noise. SNR loss scores were averaged over three lists to obtain the final SNR loss score.

Flanker and QuickSIN scores may be correlated across subjects; this was measured with a linear model for each age group, using R (R Core Team 2017).

### Stimuli and MEG recording

The task and stimuli were the same as the ones described in the previous study (Presacco et al. 2016a, 2016b). For each subject, the MEG response was recorded with a 157 axial gradiometer whole head MEG system (KIT, Kanazawa, Japan) inside a magnetically shielded room (Vacuumschmelze GmbH & Co. KG, Hanau, Germany) at the University of Maryland, College Park, sampled at 1000 Hz with online low-pass filter of cut-off frequency at 200 Hz. The stimulus was continuous speech (a narrated audio book), either from a solo speaker or a mixture of two concurrent speakers. The solo-speaker speech stimuli were one-minute segments from an audiobook, *The Legend of Sleepy Hallow* by Washington Irving, narrated by a male speaker (http://www.audiobooktreasury.com/legend-of-sleepy-hollow/). The mixture was composed of foreground speech to which the subject was instructed to attend and a background, which served as a distractor. The foreground speech was from the same source as the clean speech condition. The background stimuli were one-minute segments from an audiobook, *A Christmas Carol* by Charles Dickens, narrated by a female speaker http://www.audiobooktreasury.com/a-christmas-carol-by-charles-dickens-free-audio-book/). The foreground and background speech segments were mixed together at four different power ratios, of 3 dB, 0 dB, -3 dB and -6 dB. The foreground speech stimulus used in the -6 dB condition and the clean speech were identical, and the clean speech stimulus was only presented after all mixed speech stimuli had been presented. Stimuli were delivered through E-A-RLINK earphones inserted into the ear canal, at a comfortably loud listening level of approximately 70 dB SPL.

For each subject, under each condition, the raw MEG recording was first denoised by time-shifted principle component analysis (TSPCA; de Cheveigné and Simon 2008), in which three separate reference channels recording the environmental noise serve as a reference with which to eliminate environmental noise from the 157 neural data channels. Based on the output signal from TSPCA, a blind source separation approach, denoising source separation (DSS; de Cheveigné and Simon 2008; Särelä and Valpola 2005) was then used to estimate dominant auditory components. Based on the 2-8 Hz band-passed response (Ding and Simon 2013), DSS-based spatial filters were extracted and applied to the original signals, thus creating the DSS components which were additionally band-pass filtered between 1-8 Hz (Ding and Simon 2012a). Finally, the first DSS component was analyzed further as described below.

### Data analysis

#### Temporal mutual information function (TMIF)

To decode cortical phase-locked response to speech, a method based on mutual information was developed, based on the temporal mutual information function (TMIF). It is a non-linear analog of temporal response function (TRF) (Ding and Simon, 2012). A typical TRF has prominent peaks at latencies of approximately 50 ms and 100 ms (with opposite polarities), meaning that any speech envelope feature evokes a pair of opposite cortical responses 50 ms and 100 ms later. Since this implies enhanced cortical processing of speech information at those latencies, we may expect an enhanced level of mutual information at similar latencies (though both peaks would be positive since mutual information is nonnegative). Only the TMIF of the first DSS component is computed here.

While mutual information can naturally be applied to continuous random variables, when used in practical data analysis the continuous values are typically binned, meaning that the stimulus and response are quantized into discrete random variables. The mutual information between a stimulus *X* and a response *Y* is defined using their probability distributions. To estimate the TMIF, we first quantize both the speech envelope and the response level into 8 bins based on the equipartition principle, where the number of samples assigned to each bin is approximately the same (limited necessarily by the divisibility of the number of samples into the number of bins). Here, we denote *x*(*t*) and *y*(*t*) as the quantized speech envelope and response level at time *t*, respectively. The TMIF level at time-step *t* is defined to be mutual information between stimulus and response shifted forward by *t*,

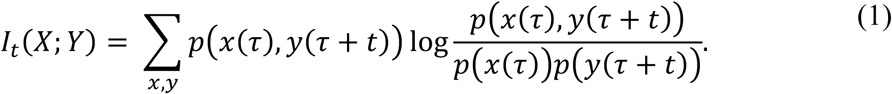

Let *S* = {1, 2, …, 8} be a set of bins from which the sample values are drawn. The joint probability distribution of *x*(*τ*)∈ *S* and *y*(*τ* + *t*)∈ *S*, i.e., *p*(*x*(*τ*), *y*(*τ* + *t*)), is drawn from different values of *τ*, which ranges from 0 to *L* − 1, where *L* is the length of the stimulus (or response) window in ms. Since the computation is at a sampling rate of 1 kHz (1 ms sampling period), *L* is also the sample size. In practice, the mutual information at each time point is estimated from its relation to entropy and conditional entropy, *I*(*X*; *Y*)=*H*(*Y*)−*H*(*Y*|*X*). With this, the equation for mutual information at a given latency *t* can be rewritten as

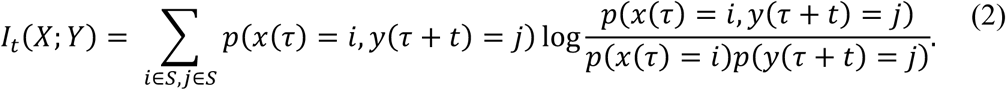

Here, *i* and *j* are values drawn from set *S*; *t* and *τ* are even integer numbers of ms, since we use a time window of 500 ms for *t* and estimate mutual information per 2-ms step, i.e., *t* ∈ {0, 2, …, 498 (*ms*)}. We then denote the TMIF function by *TMIF*(*t*) = *I*_*t*_(*X*; *Y*). In summary, *TMIF*(*t*) estimates the mutual information between the stimulus, and the response shifted forward by time *t*. If we denote *Y*_*t*_ as the response shifted forward by *t, TMIF(t)* = *I*(*X*; *Y*_*t*_); in this sense *TMIF*(*t*), the mutual information for any specified latency *t*, still relies on the entire stimulus. and entire response, as illustrated in Figure 1.

**Figure 1.**
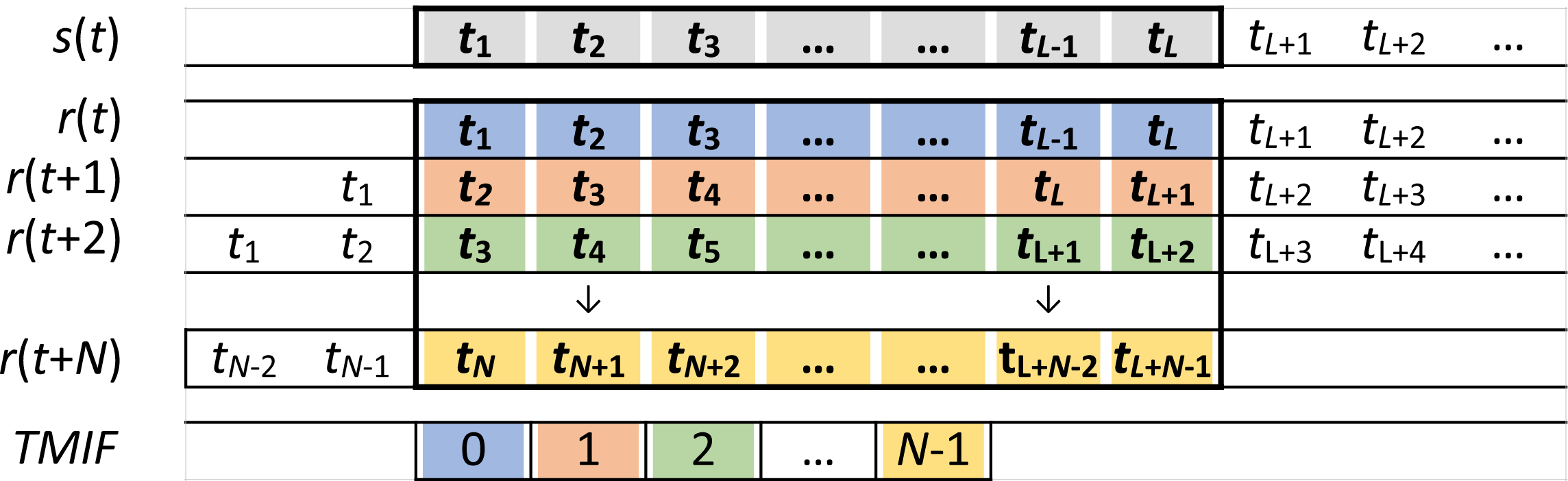
Cartoon illustration of how the TMIF is calculated for its different latencies. For *TMIF*(0), the value of the TMIF at the initial time sample (zero latency), the time-matched distributions of the entire stimulus (*s*, in gray) and the entire response (*r*, in blue) are used. For *TMIF* (1), the value of the TMIF at the next time sample, the distribution of the entire stimulus (*s*, in gray) is still used, but the delayed-by-1-sample response (*r*, in orange) is used instead of the non-delayed response. Thus, each latency value of the TMIF is computed using the entire distribution of the stimulus and the entire distribution of the appropriately delayed response.

To prove that *TMIF*(*t*) does not contain redundant information introduced by repeatedly shifting *Y*, we show that *I*(*X*; *Y*_*t*_, *Y*_*t*+1_, …, *Y* _*t*+*N*_)− *I*(*X*; *Y* _*t*+1_, …, *Y* _*t*+*N*_) = *I*(*X*; *Y*_*t*_), where *N* = 498 (*ms*). Based on the chain rule for mutual information (Cover and Thomas 1991), we have,

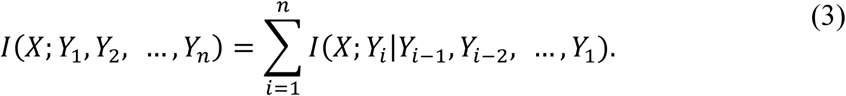

Therefore,

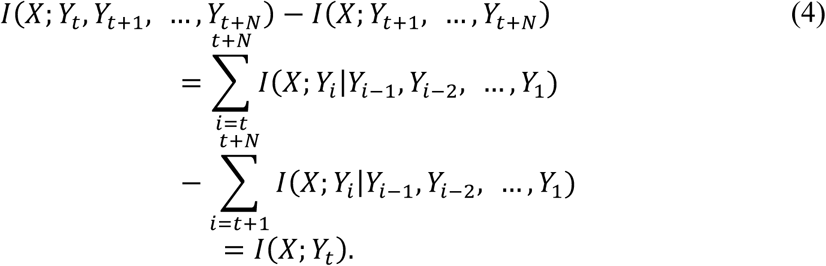

Thus *TMIF*(*t*) is not affected by repeatedly computing the mutual information from the shifted response *Y*.

After estimating the TMIF for each condition and subject, the distinctive peaks with approximate latencies at 50, 100, and 200 ms are identified as the MI50, MI100 and MI200 peaks. Peaks are found by searching for the maximum value over a specific time range. Since the response latencies differ when in quiet condition and noise conditions, different ranges are applied for different conditions, with range boundaries determined by the trough latencies in the relevant TMIF when averaged over subjects. Specifically, for the quiet condition, the MI50 corresponds to the time point with the largest amplitude in the range 2-86 ms, while the MI100 and MI200 each correspond to the maximum of ranges of 80-160 ms and 150-300 ms respectively. The group difference is tested for each peak by performing 2-sample one-tailed *t*-tests over amplitudes. For each of the noise conditions, the TMIF is analyzed analogously. TMIFs are computed for both foreground and background speech. The specific temporal ranges used for foreground TMIFs were 2-70 ms for the MI50, 50-200 ms for the MI100 and 200-300 ms for the MI200. The specific temporal ranges used for background TMIFs were 2-120 ms for the MI50, 120-230 ms for the MI100 and 200-350 ms for the MI200. The group difference is tested for each peak by performing the same *t-*tests over the averaged amplitude across SNRs.

#### Lateralization analysis

To investigate cortical lateralization, the MEG recordings were divided into two sets based on the x-coordinates (medial-lateral dimension) of the corresponding sensors in a 2-D topography (Figure 9C). DSS components were separately computed for left 79 sensors and right 78 sensors. The first DSS components for left and right sensors are representations of auditory responses for left and right hemispheres, respectively. TMIFs were estimated separately for left and right hemispheres. The HAROLD model suggests reduced lateralization for older listeners in domains of episodic memory, working memory, attention and inhibitory control (Cabeza 2002).

#### Statistics

To systematically examine relationships among neural responses properties of the TMIF (specifically the MI50, MI100 and MI200 peaks) and behavioral scores, linear mixed effect models (LME) were used. For each neural response peak, a base model was constructed as a function of fixed effects from *age* × *attention* × *behavior* + *SNR* and random effects of subject-specific bias. Here, *attention* is either foreground or background, and *behavior* is either the Flanker or QuickSIN score. The 4-way interaction was not included due to the limited degrees of freedom. To investigate the significance of a specific factor (or an interaction) in the prediction of a neural response, a second model was constructed without that factor (or interaction) and was compared with the base model by ANOVA. Then non-significant factors or interactions were excluded from model, and the significant interaction was examined by dissecting it into all possible combinations of its categorical values and further analyzed by linear models. All linear model analysis was done in R. Outlier data samples, which would have otherwise violated parametric assumptions for linear model testing (skewness, kurtosis and homoscedasticity), were detected and excluded using *gvlma* in R (Peña and Slate 2006). LME analysis was done by the toolbox *lme4* (Bates et al. 2015), and the linear model without random effects was analyzed using the *lm* function in R (Chambers 1992; Wilkinson and Rogers 1973). A stepwise regression test was performed in SPSS to test for linear contributions of Flanker score and MI200 level to speech intelligibility. Where appropriate, *t* tests for significance were supplemented with effect size (Cohen’s d) and its 95% confidence interval (CI). When the CI excludes zero, this is alternate evidence that the result is statistically significant (i.e., the effect size is significantly greater than zero at an *α* level of 0.05).

## Results

By implementing the approaches established above, for each subject under each condition, TMIFs were computed for the first DSS component. Here, we report results under the conditions of clean speech and mixed speech with SNRs of +3, 0, -3 and -6 dB, and lateralization analysis.

### Behavioral correlation

A linear model of *QuickSIN* ∼ *Flanker* was examined, separately for younger and older listeners, to test the relationship between Flanker score and QuickSIN score. The assumptions for linear models of skewness, kurtosis and homoscedasticity were all satisfied (using *gvlma* in R). Results show a significantly negative regression slope for older listeners (*t*_13_ = −2.21, *p* = 0.046), but not for younger listeners (*t*_15_ = 0.16, *p* = 0.873). Linear model assumption testing for the older listeners showed a low kurtosis value, 0.09 (*p* = 0.767), avoiding the need to treat any data points as possible outliers.

**Figure 2.**
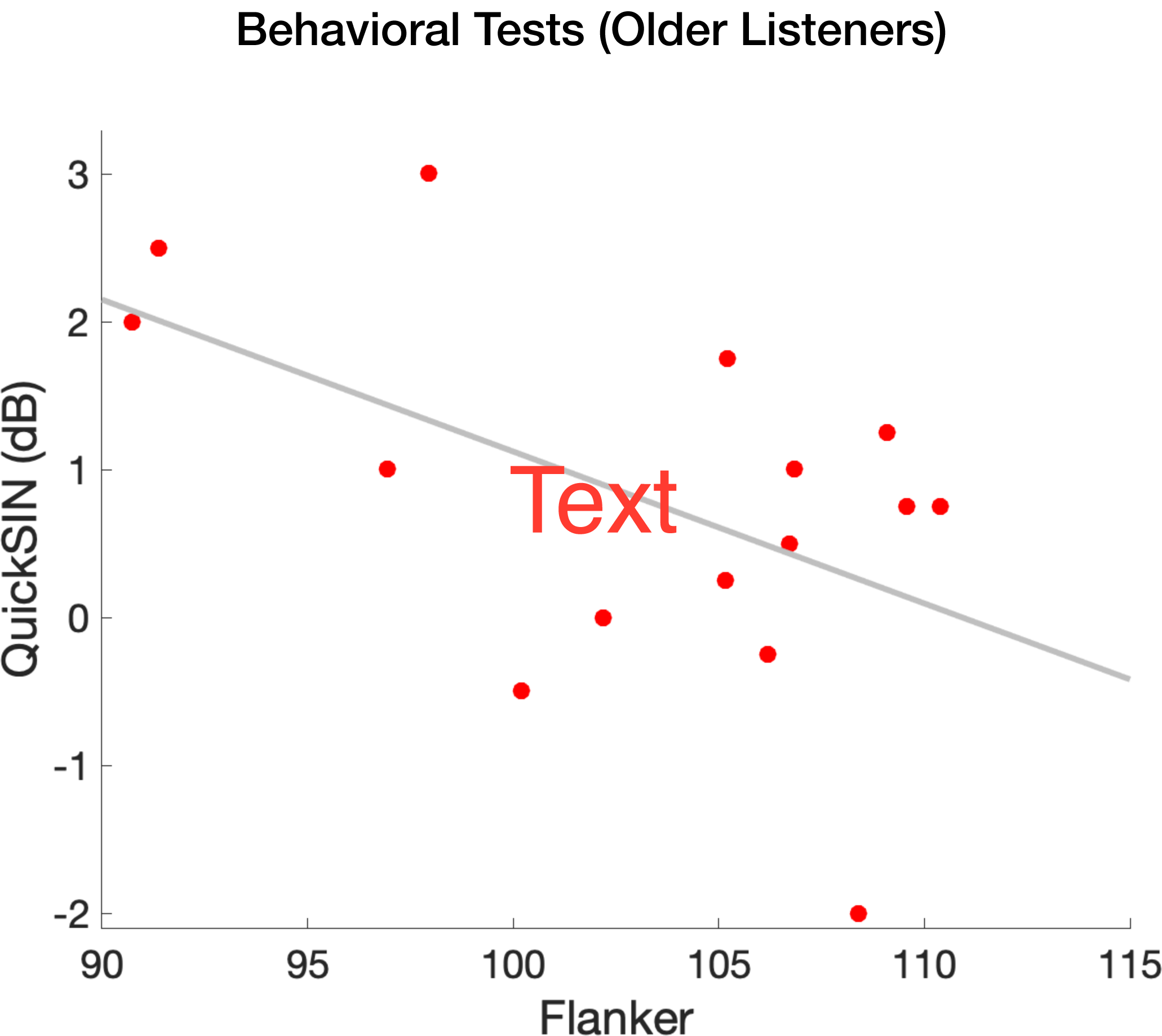
Behavioral tests. Flanker score (higher is better) is negatively correlated with Quick-SIN score (lower is better) in older listeners but not younger listeners.

### Neural Responses to Clean speech

To investigate age differences in the quiet condition, peaks analogous to TRF peaks were identified, i.e., the MI50, MI100 and MI200 (analogous to the M50, M100 and M200 MEG TRF peaks and similarly named evoked response peaks). As with their counterparts, peaks of different latencies may be associated with different stages of the processing chain. A one-tailed *t-*test was performed for each peak amplitude for younger against older. Results show that all the peaks from the older listeners are significantly larger than those of the younger (*t*_30_ = −1.85, *p* = 0.037 for MI50, *t*_30_ = −2.52, *p* = 0.009 for MI100 and *t*_30_ = −2.24, *p* = 0.031 for MI200). The results suggest that all the processing stages in the aging cortex have an exaggerated response to the clean speech envelope.

**Figure 3.**
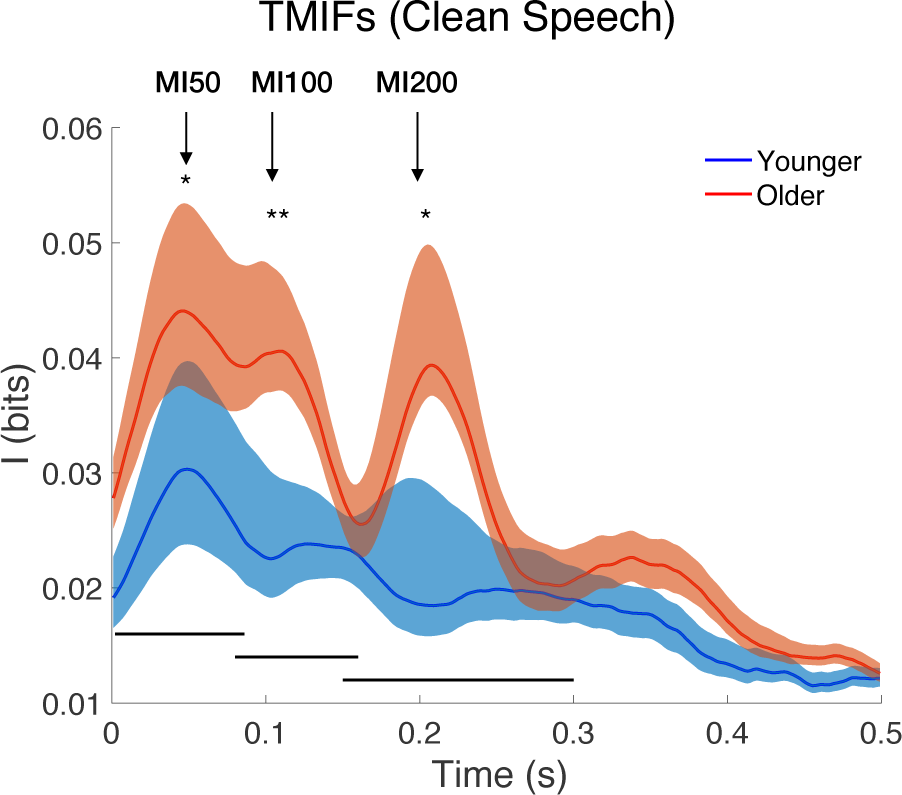
TMIF to clean speech. Shaded areas above and below the solid lines indicate the standard error of mean. The temporal ranges over which MI50, MI100 and MI200 for each subject are constrained are marked by the three black lines above x-axis. Asterisks show the significance of amplitude differences between the two groups from a one-tailed *t*-test (\**p*<0.05, \*\**p*<0.01).

### Neural Response to Mixed Speech

In mixed speech conditions, separate TMIFs for both foreground and background speech were computed, as shown in Figure 4 and Figure 5, respectively. Response peaks were extracted, and effects from factors of age, attentional focus and behavioral score were examined systematically by linear mixed effect models, *MI level*∼ *age* × *attention*× *behavior* + *SNR* + (1|*subject*). In the model, the random effects term, (1| *subject*), allows for subject-specific intercepts or bias, and *behavior* is either Flanker or QuickSIN. When *behavior* is Flanker, the 3-way interaction is significant for models predicting the amplitude of the MI50 (*χ*^2^_4_ = 16.45, *p* = 0.002), MI100 (*χ*^2^_4_ = 98.08, *p* < 0.001) and MI200 (*χ*^2^_4_= 91.38, *p* < 0.001) compared with a null model with no interactions, i.e., *MI* ∼ *age* + *attention* + *behavior* + *SNR* + (1|*subject*). To examine the significance of interactions, variables *age, attention* and *behavior* were then separately released from the 3-way interaction. Those results show that the *age* × *attention* interaction is significant in predicting the amplitude of the MI50 (*χ*^2^_3_ = 7.61, *p* = 0.055 by releasing *behavior* (*Flanker*); *χ*^2^_3_ = 14.17, *p* = 0.003 by releasing *age*; *χ*^2^_3_ = 14.52, *p* = 0.002 by releasing *attention*, and the 3-way interaction is significant in predicting the amplitude of the MI100 (*χ*^2^_3_ = 66.89, *p* < 0.001 by releasing *behavior* (*Flanker*); *χ*^2^_3_ = 70.89, *p* < 0.001 by releasing *age*; *χ*^2^_3_ = 83.92, *p* < 0.001 by releasing *attention*) and MI200 (*χ*^2^_3_ = 88.98, *p* < 0.001 by releasing *behavior* (*Flanker*); *χ*^2^_3_ = 78.67, *p* < 0.001 by releasing *age*; *χ*^2^_3_ = 72.39, *p* < 0.001 by releasing *attention*). Therefore, variables of *age* and *attention* interact with *behavior* in predicting the level of mutual information, and the prediction power changes for different combinations of *age* × *attention*, such as younger and foreground vs. older and foreground. To examine the prediction differences, the model of *MI* ∼ *behavior* + *SNR* was constructed separately for different combinations of *age* and *attention*. The overall model significances are shown in Table 1, and the effects of *behavior* are shown in Table 2.

**Table 1.**
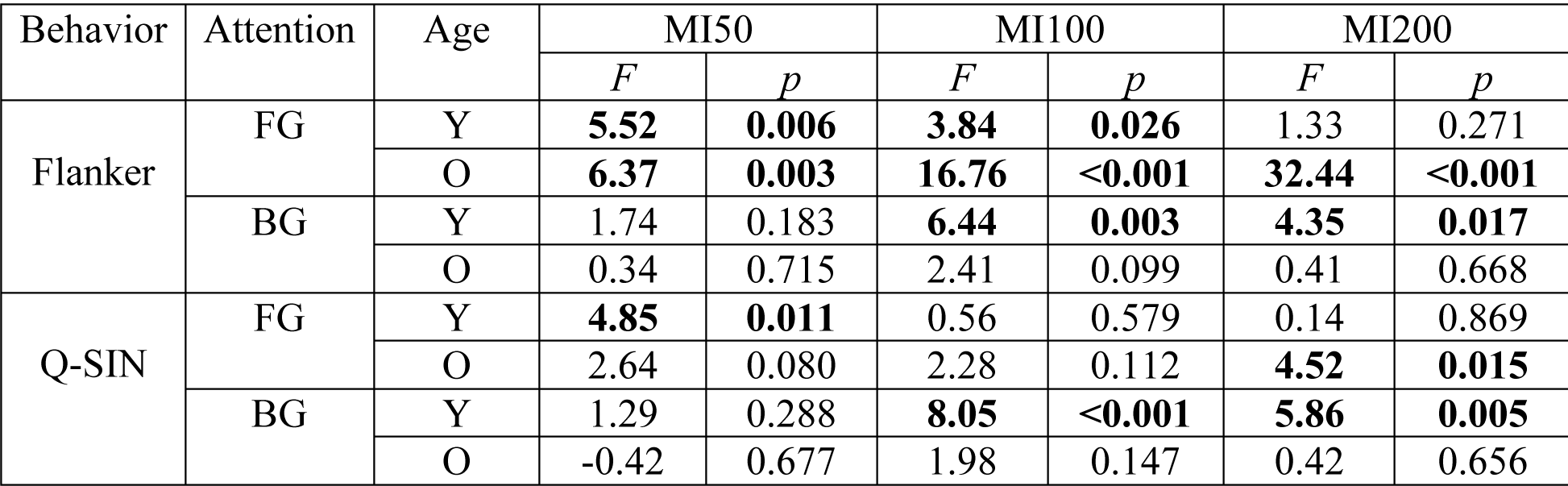
Model *MI* ∼*behavior* + *SNR* significance. FG: foreground; BG: background; Y: younger; O: older. Significant findings are in boldface.

**Table 2.**
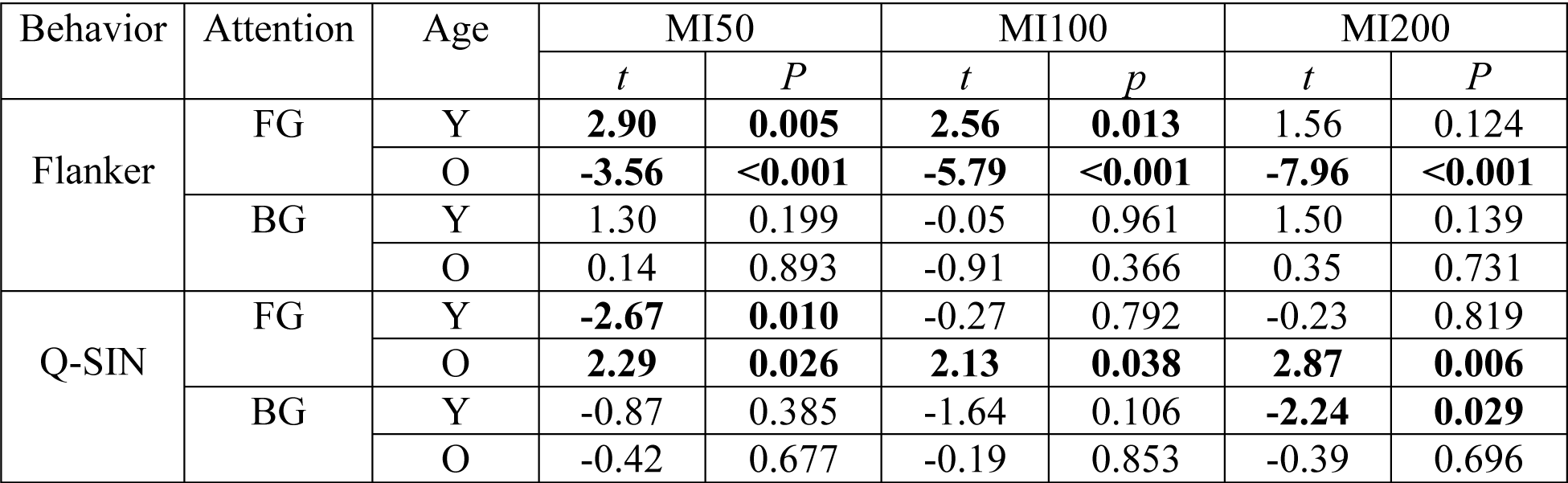
Effects of behavioral scores (Flanker and Quick-SIN) in prediction of mutual information. Significant findings are in boldface.

**Figure 4.**
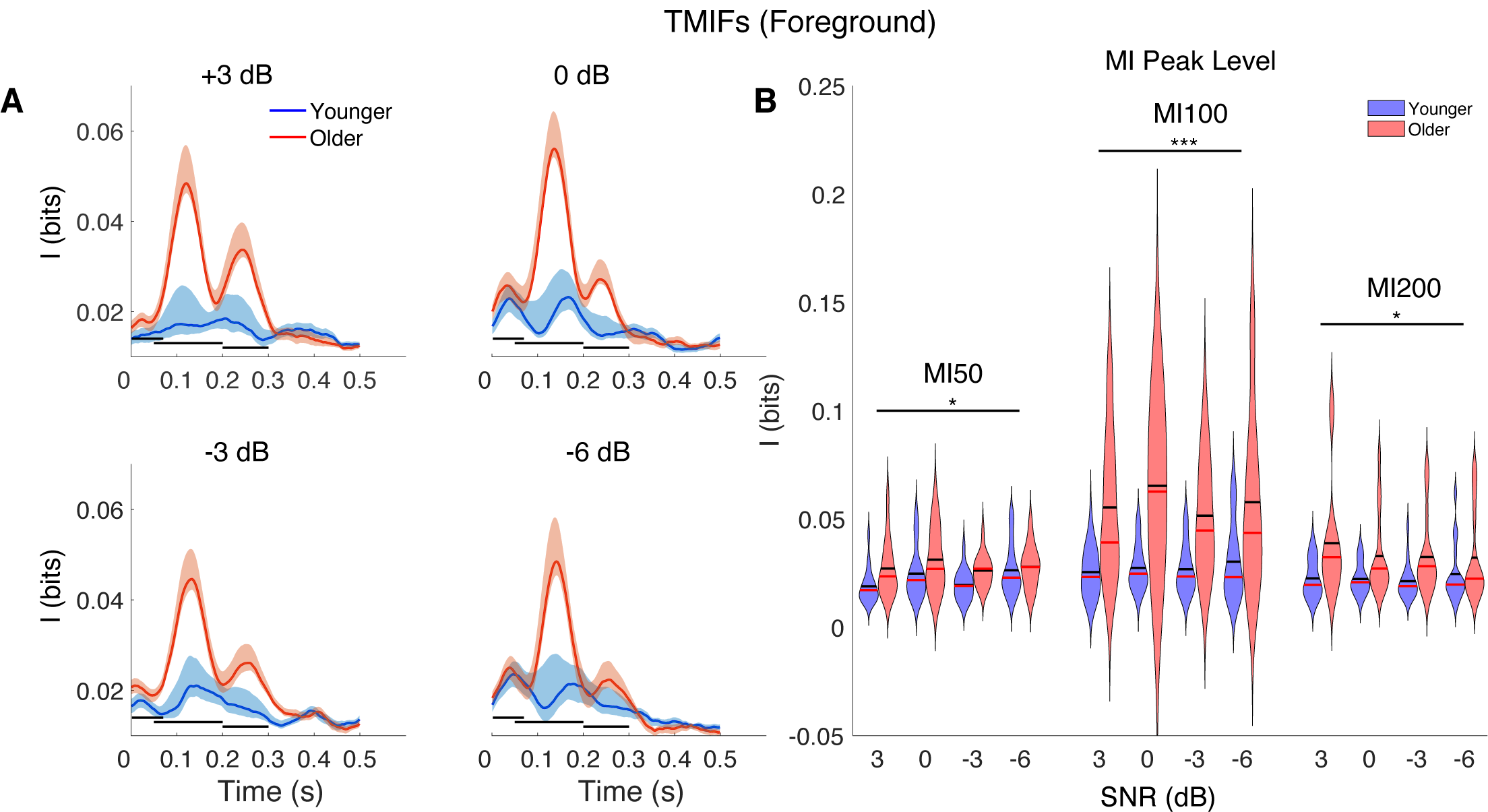
TMIFs of the foreground speech are exaggerated in older listeners. **A**. The four plots illustrate different SNR conditions of 3, 0, -3, and -6 dB SNR, with younger listeners in blue and older listeners in red. The three black horizontal lines in each figure indicates the ranges from which three peaks are extracted. Shaded areas: ± 1 SEM. **B**. MI peak level in older (red violin plots) and younger listeners (blue violin plots). 2-sample one-tailed *t*-tests on the averaged peak amplitudes over SNR conditions show that the older listeners have significantly larger amplitudes (\**p*<0.05, \*\**p*<0.01, \*\*\**p*<0.001). In each violin plot, the black bar indicates mean value, and the red bar indicates the median.

**Figure 5.**
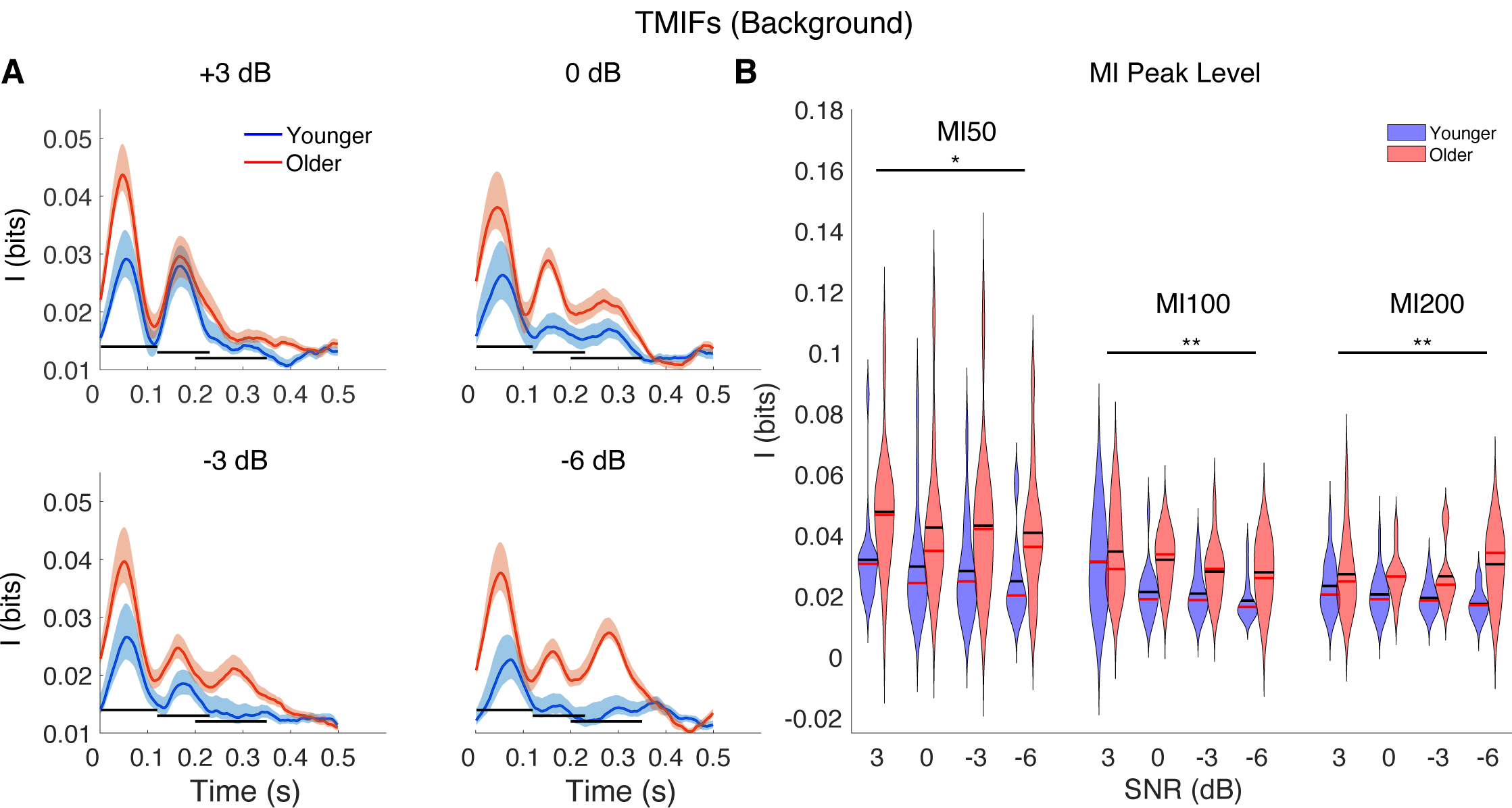
TMIFs of background speech are exaggerated in older listeners. Plots in **A** illustrate different SNR conditions of 3, 0, -3 and -6 dB, with younger listeners in light blue and older listeners in light red. The three black horizontal lines in each figure indicates the ranges from which three peaks are extracted. Shaded areas: ± 1 SEM. Figure **B** compares peak amplitudes in older listeners (red violin plots) with younger listeners (blue violin plots). Similar to the responses to foreground speech, the older listeners’ responses have significantly larger peaks than younger listeners with 2-sample one-tailed *t-*tests on the averaged peak level over SNR conditions (\**p*<0.05, \*\**p*<0.01, \*\*\**p*<0.001). Additionally, the MI50 level is notably larger than the other two peaks, for both groups. This is consistent with a representation-suppression mechanism for background processing. In each violin plot, the black bar indicates mean value, and the red bar indicates the median.

To investigate whether the age-related exaggerated response occurs for both foreground and background, and which peaks might contribute, mutual information levels of all three peaks, for each stimulus, under each SNR condition were found for each subject and compared between groups. Older listeners show significantly larger mutual information levels in all three peaks for both foreground (*t*_30_ = −2.07, *p* = 0.024 for MI50, *t*_30_ = −3.80, *p* < 0.001 for MI100 and *t*_30_ = −2.37, *p* = 0.012 for MI200) and background (*t*_30_ = −2.44, *p* = 0.010 for MI50, *t*_30_ = −2.57, *p* = 0.0076 for MI100 and *t*_30_ = −2.90, *p* = 0.0035 for MI200). Therefore, both foreground and background representations are exaggerated for older listeners, with the MI100 showing the largest effect.

### MI200 relationships with behavioral performance

As can be seen in Figures 4 and 5, the dependence of the MI200 peak level on SNR condition exhibits different trends for older and younger listeners. Notably, for younger listeners, the MI200 response remains steady as SNR decreases for foreground speech while it decreases for background speech. However, for older listeners, the response to foreground decreases as SNR decreases, while the response to background increases as SNR decreases. MI200 saliency is then defined as the difference between foreground and background information (Figure 6A, third row), and any trends as a function of SNR can be analyzed via the slope of difference-by-SNR linear regression line (Figure 6B, third row). A right-tailed 2-sample *t*-test is performed on the slopes of younger listeners against the older, resulting in a significantly larger slope for younger than older listeners (*t*_30_ = 2.31, *p* = 0.014). To test the positivity of the ratio as SNR decreases in younger participants, a right-tailed 1-sample *t-*test is conducted on the slopes of younger listeners, and the results show a significant positive trend as SNR decreases (*t*_16_ = 1.83, *p* = 0.043; *d* = 0.43, 95%*CI* = [0.20 × 10^−5^, +∞]). Similarly, a left-tailed 1-sample *t-*test against zero on slopes of older listeners show a negative trend but not significant (*t*_14_ = −1.47, *p* = 0.083; *d* = −0.34, 95%*CI* = [−∞, 0.86 × 10^−4^]) (Figure 6B). In short, age does affect the response pattern (with increasingly challenging mixed speech conditions) of this late cortical representation.

**Figure 6.**
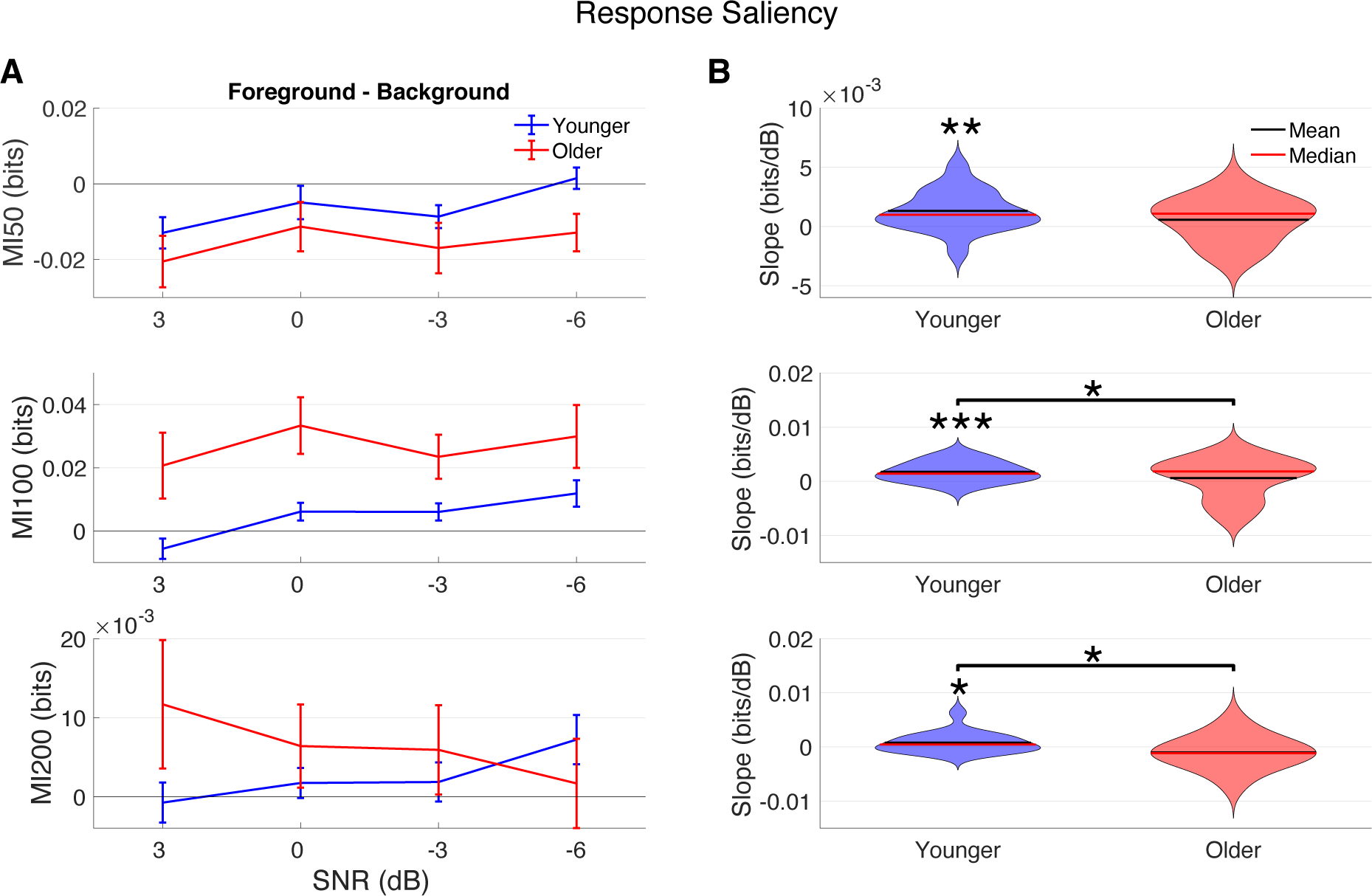
MI peak level difference between foreground and background as a function of SNR in younger and older listeners for the MI50 (upper), MI100 (middle), and MI200 (lower) peak levels (**A**) and their slopes (**B**). **A**. Younger listeners (blue) demonstrate an increasing trend with decreasing SNR for all three MI peaks, while the older (red) demonstrate a decreasing trend for the MI200 peak. **B**. MI ratio slopes as a function of SNR for individuals in the two age groups. Younger listeners have a significantly positive slope for all three MI peaks (linearly fitted regression to the data shown in panel A), while older listeners show a weakly negative slope (not statistically significant) for MI200 and weakly positive slopes for MI50 and MI100 (not statistically significant). The slope difference between groups is significant for MI100 and MI200. (\**p*<0.05, \*\**p*<0.01, \*\*\**p*<0.001)

The different MI200 saliency trend by age suggests functional differences in neural suppression of the background and/or enhancement of foreground representation for older listeners as SNR level decreases. These abilities may be related to inhibitory and attentional control. A linear model of *MI*200 ∼ *Flanker* + *SNR* was tested separately for younger and older listeners. For younger listeners, the model is significant (*F*_2,59_ = 3.28, *p* = 0.044), giving a significantly positive MI200-Flanker slope (*t*_59_ = 2.26, *p* = 0.028), but with no effect of SNR (*t*_59_ = 1.05, *p* = 0.300). For older listeners, the model shows an even stronger effect size (*F*_2,54_ = 40.29, *p* < 0.001) and a significantly negative MI200-Flanker slope (*t*_54_ = −8.97, *p* < 0.001); however, no significant effect of SNR is observed (*t*_54_ = 0.79, *p* = 0.431). Additionally, a separate linear model of *MI*200 ∼ *Flanker* under each SNR level was tested. Linear model assumptions were satisfied in each test. For younger listeners, the MI200-Flanker models are not significant (*t*_15_ = −0.11, *p* = 0.917 for +3 dB, *t*_15_ = 0.31, *p* = 0.764 for 0 dB, *t*_15_ = 0.79, *p* = 0.443 for -3 dB, *t*_15_ = 1.70, *p* = 0.109 for -6 dB). However, for older listeners, MI200-Flanker slope is significantly negative in all SNRs (*t*_15_ = −2.28, *p* = 0.040 for +3 dB, *t*_13_ = −4.42, *p* < 0.001 for 0 dB, *t*_13_ = −5.19, *p* < 0.001 for -3 dB, *t*_13_ = −6.91, *p* < 0.001 for -6 dB). Example scatter plots are shown in Figure 7.

**Figure 7.**
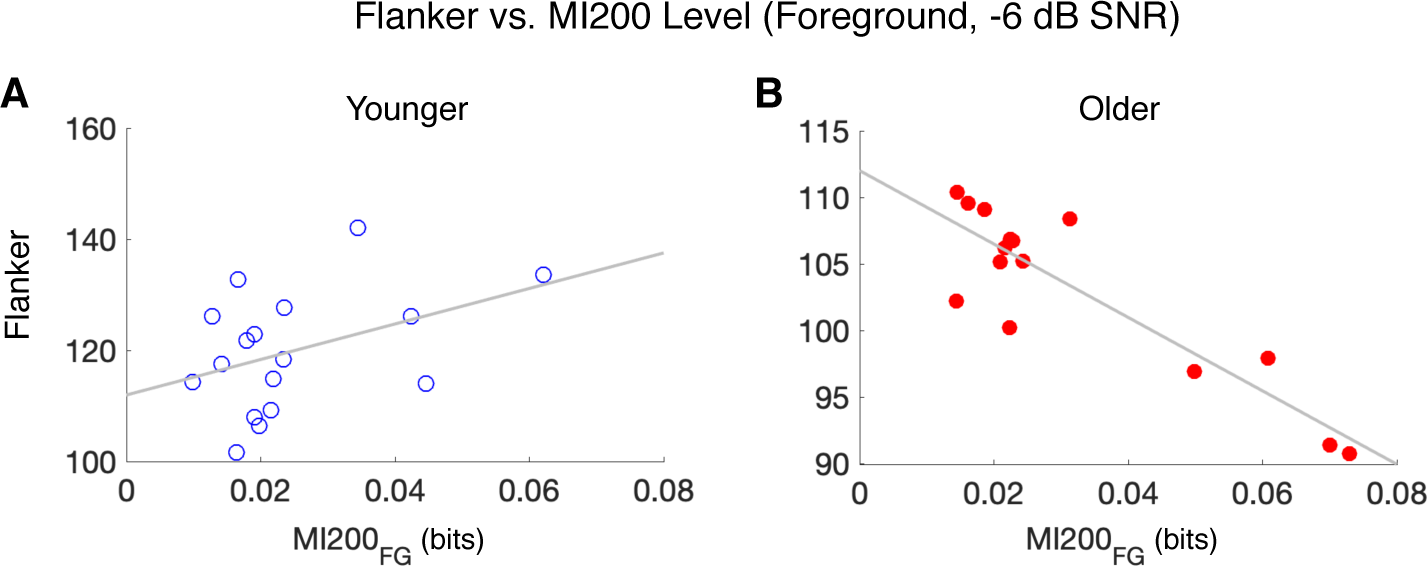
Relationship between foreground MI200 level and Flanker test score by age for a difficult SNR. Scatterplots of foreground MI200 level and Flanker test scores under the most challenging condition, -6 dB SNR, for younger listeners in **A** (blue) and older listeners in **B** (red). Linear regression lines in gray were determined by the corresponding linear models.

Since the speech-in-noise behavioral score is negatively associated with the Flanker inhibition score in older listeners (Figure 2), the foreground MI200 level might also be associated with the QuickSIN score. A stepwise regression (backward method) testing for linear contributions of Flanker score and MI200 level to QuickSIN score shows that only MI200 level, but not Flanker score, contributes to QuickSIN level (*F*_1,13_ = 7.27, *p* = 0.018; third sub-table in Table 3). Full model results are shown in Table 3. These results demonstrate that higher MI200 level is associated with worse speech-in-noise performance for older listeners. Scatter plots are shown in Figure 8.

**Table 3.**
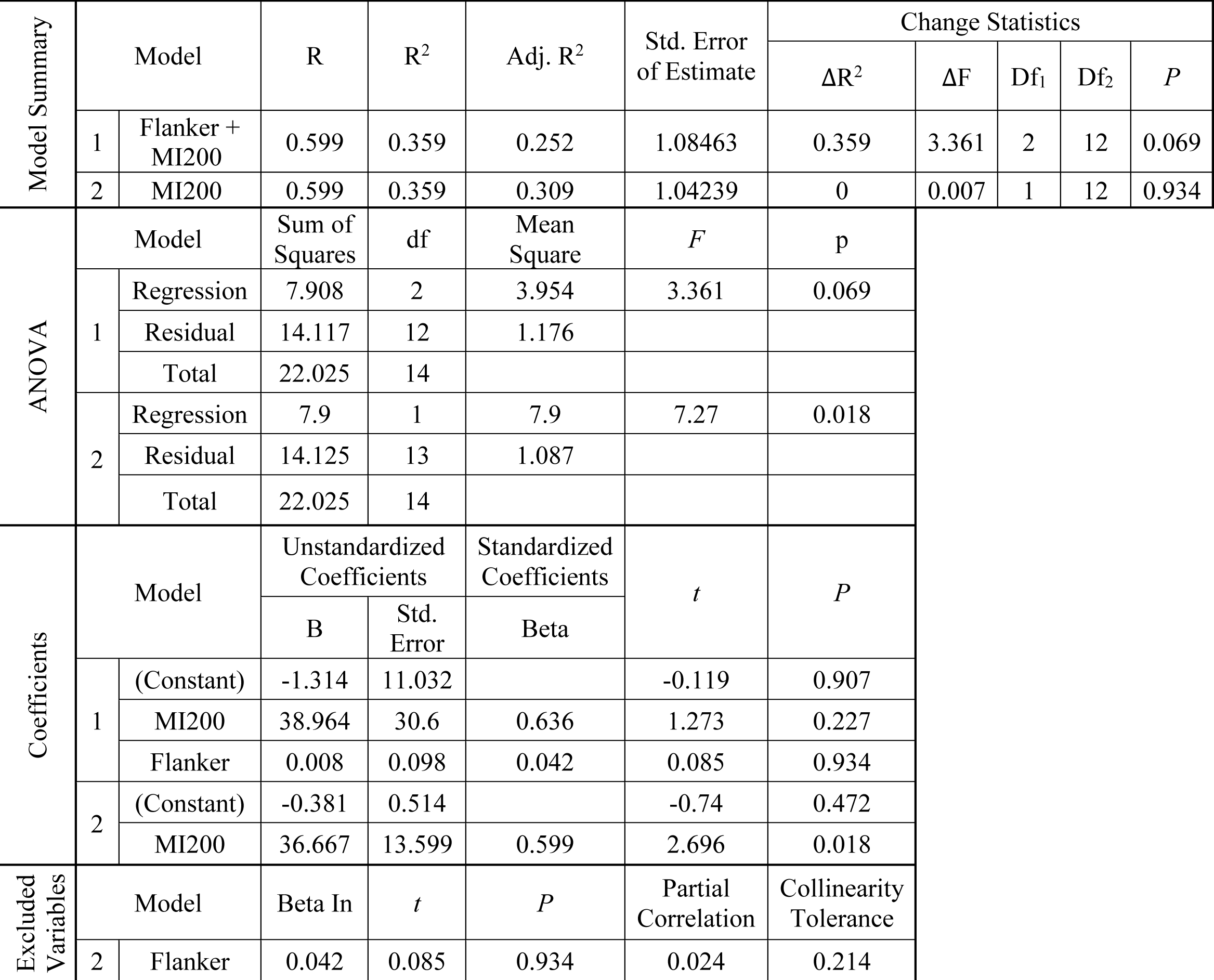
Stepwise regression of *QuickSIN* ∼ *Flanker* + *MI*200 for older listeners (backward method). The model summary introduces the full (model 1) and reduced (model 2) models: 1) QuickSIN modeled as dependent on both Flanker score and MI200 level; 2) the same model but with Flanker score selected as an excluded independent variable. ANOVA results show that only the second model is significant. The overall results suggest that only MI200 level, but not Flanker score, predicts QuickSIN score.

**Figure 8.**
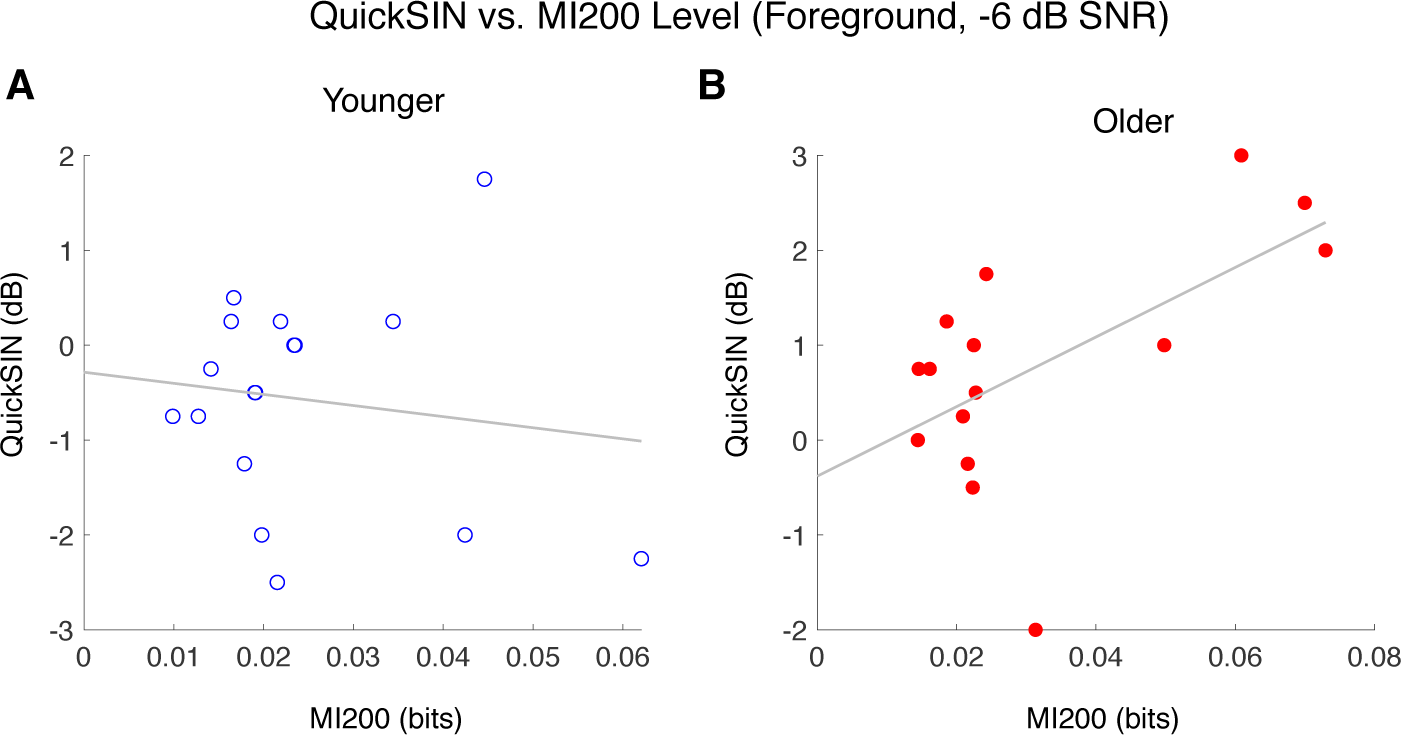
Scatter plots of foreground MI200 level and speech-in-noise performance. **A**. No significant association is seen for younger listeners. **B**. The association is significant in older listeners (red). Stepwise regression analysis shows only MI200 level but not Flanker contributes to predicting QuickSIN performance. Linear regression lines in gray for both plots were determined by the corresponding linear models.

**Figure 9.**
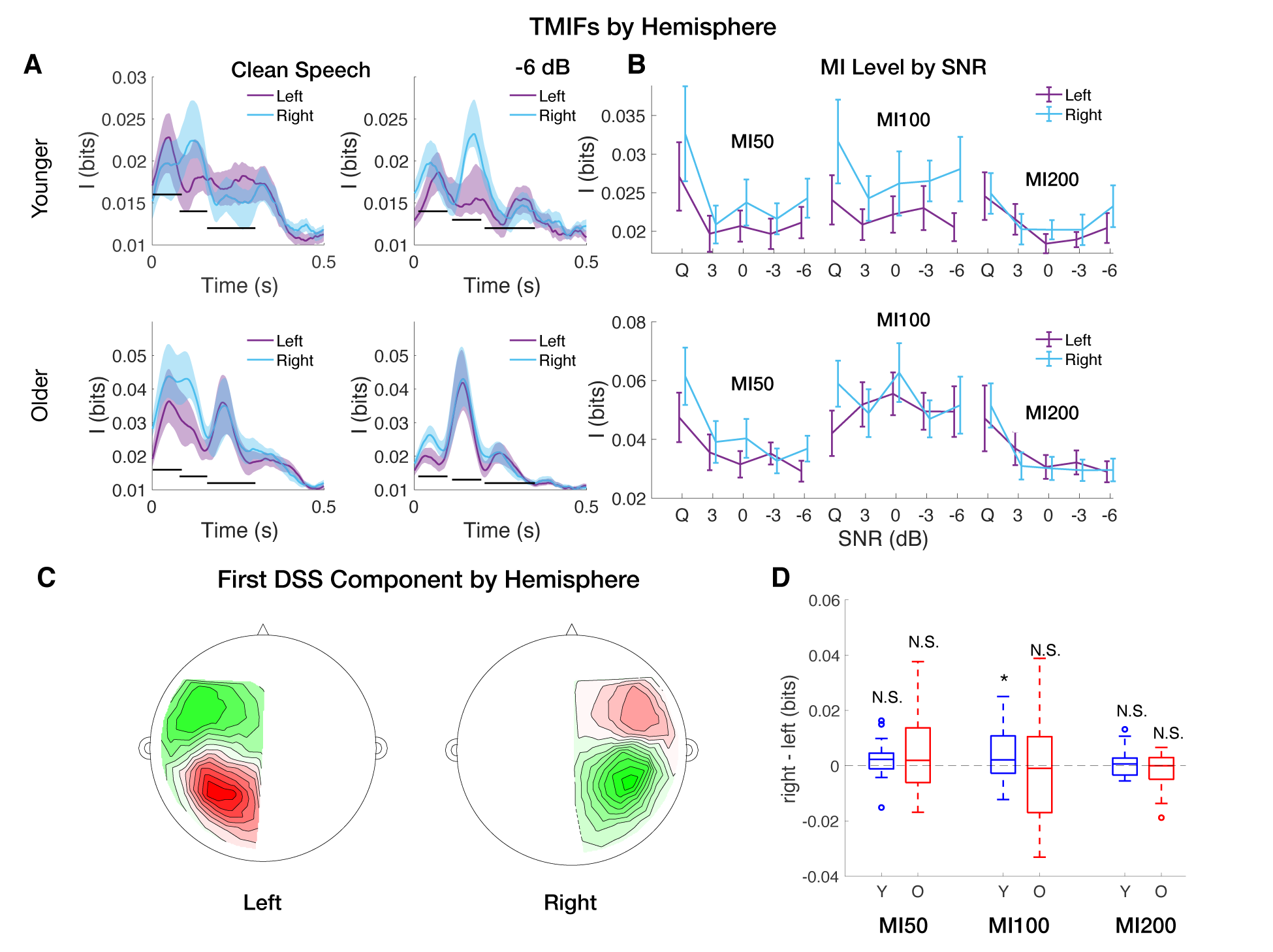
Lateralization analysis. **A**. TMIFs by hemisphere for younger (first row) and older (second row) listeners under clean speech and -6 dB conditions, with left hemisphere in purple and right hemisphere in light blue. **B**. MI50, MI100 and MI200 trends by conditions. The x-axis labels condition by SNR, or where ‘Q’ (quiet) is the clean speech condition. **C**. Topographies of the first DSS component for left and right hemispheres for an example subject. **D**. The difference between right and left hemispheres in MI levels averaged across SNRs. The difference for younger listeners is significantly larger than zero for the MI100, however, no difference is seen for older listeners.

### Lateralization

TMIFs are estimated for both left and right hemispheres, and the difference between hemispheres were examined for all three peak levels (Figure 9). A linear model of *MI level* (*right*)− *left*)∼ *age* × *SNR* was tested with *lm* in R, separately for each peak. For the MI50, results indicate that the model is not significant (*F*_3,124_ = 0.22, *p* = 0.885). A one-tailed *t*-test on the right-left difference of MI50, averaged across SNRs, against zero, shows that MI50 level difference is not significantly larger than zero for both younger listeners (*t*_16_ = 1.26, *p* = 0.112; *d* = 0.31, 95%*CI* = [−0.28 × 10^−3^, +∞]) and older listeners (*t*_14_ = 1.51, *p* = 0.077; *d* = 0.39, 95%*CI* = [−0.27 × 10^−4^, +∞]). For the MI100, results also indicate that the model is not significant (*F*_3,124_ = 0.44, *p* = 0.725). In this case, a one-tailed *t-*test shows that the MI level difference for younger listeners is significantly larger than zero, (*t*_16_ = 1.89, *p* = 0.038; *d* = 0.46, 95%*CI* = [0.98 × 10^−4^, +∞]), but not for older listeners (*t*_14_ = 0.77, *p* = 0.229; *d* = 0.2, 95%*CI* = [−0.0014, +∞]), suggesting a right-lateralized response for younger and a bilateral response for older. For the MI200, however, the linear model is statistically significant (*F*_3,124_ = 2.83, *p* = 0.041) and significantly affected by age (*t*_124_ = 2.04, *p* = 0.044) with an average group difference of 0.0035 bits. This suggests that the MI200 response is more right-lateralized for younger listeners than older. However, one-tailed *t-*tests for both younger and older listeners show no lateralization for either younger listeners (*t*_16_ = 0.66, *p* = 0.259; *d* = 0.16, 95%*CI* = [−0.0016, +∞]) or older (*t*_16_ = −0.286, 2 = 0.610; *d* = 0. −0.07, 95%*CI* = [−0.002 + ∞]), indicating a bilateral MI200 response for both groups (though with a greater right-hemisphere bias for younger listeners).

## Discussion

### Mutual information vs. linear methods

By developing a novel approach based on information theory, phase-locked cortical responses to the speech envelope can be measured without resorting to linear-only statistics. The TMIF unveils different processing stages in the cortical response to speech, via the mutual information peaks MI50, MI100 and MI200. Previous analysis restricted to linear methods has been done on this same dataset using both TRF analysis and stimulus reconstruction analysis. TRF analysis was able to find a group difference only for the earlier response peak, the M50 (Brodbeck et al. 2018), while the present mutual information analysis shows that all three of these peaks are significantly larger for older adults than younger adults. The group difference seen here for the MI100 and MI200 demonstrates a statistical advantage for mutual information over TRF analysis. Additionally, the late response, MI200, differs in its profile from the earlier components, in that the difference between foreground and background levels has a different pattern of dependencies on SNR for the two age groups: while the ratio in younger listeners increases with worsening SNR, it decreases in older listeners. Earlier analysis using stimulus reconstruction was able to show that foreground stimulus reconstruction accuracy is negatively correlated with Flanker score (Presacco et al. 2016b), but, critically, only when integrated over all latencies and averaged across SNR levels. The results here are far more specific: mutual information analysis shows: a) it is specifically the late response, the MI200, that negatively correlates with Flanker inhibition scores, and regardless of SNR level (Figure 7); and b) that MI200 level is also correlated with QuickSIN even after accounting for associations between QuickSIN and Flanker scores (Table 3 and Figure 8). Therefore, compared with linear methods, the analysis based on mutual information has greater statistical power in detection of group differences, and relationships between neural representation and behavioral scores.

Why this non-linear, information-theoretic analysis technique would outperform the more standard linear analysis techniques is an open question. It may be that using linear-only methods ignores critical non-linearities in the neural responses, and that those non-linearities are particularly well captured by this measure. Another possibility is that some areas of auditory cortex are actually tuned, computationally, to maximize the mutual information between the stimulus and their responses.

### Correlation between auditory and visual behaviors for older listeners

Our results show a correlation between the QuickSIN score and the Flanker visual inhibitory score for older listeners but not for younger listeners (Figure 2). Previous studies report a decline in cognitive functions including attention, visual information processing, working memory and episodic memory for older adults (Craik and Salthouse 2000; Ebaid and Crewther 2019). According to the “inhibitory deficit hypothesis”, the decline in cognitive functions are associated with an across-modality inability to reduce interference from task-irrelevant information (Hasher 2015; Hasher and Zacks 1988), and such inability presents in both auditory processing (Stothart and Kazanina 2016) and visual processing (Gazzaley et al. 2008). Our results suggest that the inability to reduce interference in both auditory and visual systems may share a common neural origin.

### Exaggerated response in the aging cortex: potential mechanisms

An exaggerated speech cortical representation for older listeners is seen at every latency considered (MI50, MI100 and MI200) and in both clean speech and adverse conditions. The age-related exaggeration in MI50 and MI100 is consistent with previous findings in auditory cortical evoked responses. The early cortical evoked P1 response (∼50 ms) has been seen to show an exaggerated response in older listeners (Woods and Clayworth 1986; Roque et al. 2019). Studies on auditory gap detection also show a larger P1 for older listeners than younger (Lister et al. 2011; Ross et al. 2010), suggesting altered neural inhibition may be responsible for this increase in amplitude. Larger N1 (∼100 ms) responses in older listeners have also been seen (Chao and Knight 1997), with Anderer et al. (1996) showing the N1 amplitude increasing linearly with age. Rufener et al. (2014) also found a larger N1 amplitude for older listeners in response to both speech and non-speech stimuli in selective attention tasks. This exaggerated response might be associated with task-related cognitive effort based on a tone classification task (Rao et al. 2010), where N1and P1 are enhanced during more difficult noise classification. However, P2 (∼200 ms) responses to tones and gaps in noise, interestingly, do not show increased amplitude for older listeners (Alain and Snyder 2008; Lister et al. 2011). This might indicate speech processing shares less with tone processing at those longer latencies. All these age-related increases in auditory ERP amplitude may be related to impaired inhibitory functions along the afferent and efferent auditory pathways (Alain and Woods 1999; Chao and Knight 1997); the aging auditory cortex shows more difficulty filtering out task-irrelevant stimuli and may require more cortical resources to process the same information (Alain et al. 2004; Pichora-Fuller et al. 2017).

Several possible mechanisms might underlie these findings. One possible contribution to exaggerated cortical representations may be a loss of neural inhibition (Caspary et al. 2008; Takesian et al. 2012). Animal studies show decreased release of inhibitory neurotransmitters, such as gamma-aminobutyric acid (GABA), in auditory cortex (Juarez-Salinas et al. 2010; de Villers-Sidani et al. 2010). Such a reduction in neural inhibition might occur as part of a compensatory gain mechanism (Caspary et al. 2008; Takesian et al. 2012), and may have broad consequences (Recanzone 2018). The aging midbrain shows deficits in temporal processing acuity in normal-hearing CBA mice (Walton et al. 1998), and the cortex is able to restore auditory processing even with a cochlear denervation and virtually eliminated brainstem response (Chambers et al. 2016). Similar exaggerated responses are also seen in cases of tinnitus and hyperacusis, at multiple levels along the auditory pathway (Auerbach et al. 2014). Since the loss of neural inhibition occurs in subcortical and cortical structures of auditory system, it may lead to an exaggeration in neural activity regardless of response latency.

Another potential contributor to exaggerated response in the aging cortex might be the utilization of more neural resources in cognitive processing, such as redundant local processing (Peelle et al. 2010) or enhanced attention (Presacco et al. 2016a). Older listeners allocate more neural resources outside the core sentence-processing network and demonstrate reduced coherence between activated regions (Peelle et al. 2010), which might, in turn, cause neighboring cortical sources to process same stimulus information independently, and thus leading to an over-representation (Presacco et al. 2016b). This effect might contribute to an exaggerated representation in any of the three peaks. Enhanced attention, in contrast, would most likely be reflected in the response with latency ∼100 ms (Ding and Simon 2012a, 2012b), which then could contribute to a larger MI100 for older listeners.

Additionally, cortical representations enhanced by additional contextual information in older listeners might also contribute to an exaggerated level of mutual information. Older listeners’ speech understanding benefits from different levels of supportive context, at sentential, lexical, phonological and sub-phonemic levels (Pichora-Fuller 2008). Embedded within the frequency range of 1-8 Hz (Cogan and Poeppel 2011), such contextual information enhancement for older listeners may be reflected by an exaggerated MI level at late latency, MI200, which is late enough to benefit from such high level information.

### Long latency processing, distractor suppression and speech-in-noise intelligibility

For these reasons, the MI200, the latest of the three components, is a viable candidate for reflecting an extra stage of speech processing that makes additional use of redundant speech information. The negative correlation between the MI200 and the Flanker score suggests that this later neural activity might serve as a bio-marker for degraded behavioral inhibitory control for older listeners. The finding is also consistent with a recent study where worse cognitive scores were found to be associated with enhanced envelope tracking (Decruy et al. 2019). The current results also show that a worsened exaggerated MI200 at the most challenging noise condition is associated with worse speech understanding. This relationship suggests that the exaggerated response, though perhaps compensatory, may not be not beneficial (or not beneficial enough) for older listeners. This might arise from an imbalance between neural excitatory and inhibitory mechanisms (Caspary et al. 2008). Alternatively, the exaggerated neural representation might be associated with a compensatory mechanism, where additional cortical regions are engaged to accomplish a difficult listening task (Presacco et al. 2016a, 2016b; Takesian et al. 2012; Wong et al. 2010). Notice that for older listeners, the MI200 peak level decreases with worsening SNR, possibly because the response to background grows stronger as SNR decreases. This suggests that even with compensatory processing, older listeners may still fail to suppress the representation of the background speech as it reaches higher sound levels. Older listeners show a trend, as SNR decreases, for MI200 saliency (foreground over background) that is consistent with this hypothesis. The MI200 saliency for younger listeners, however, for whom these SNRs cause only modest difficulty, show a slope in the direction opposite to this hypothesis. Finally, note that Decruy et al. (2019) find that enhanced envelope tracking is positively correlated with speech understanding, not negatively, but using a measure that incorporates all latencies, not just the MI200.

### Lateralization of auditory processing

In cocktail party scenarios, the MI100 shows a bilateral response for older listeners, in contrast to a right-lateralized response for younger listeners. The asymmetric neural representations for younger listeners support the ‘asymmetric sampling in time’ hypothesis for auditory processing (Poeppel 2003), where right hemisphere extracts speech information from long integration windows (∼150-250 ms). The tendency towards neural activity symmetry with aging is consistent with the HAROLD model, where memory, attention and inhibitory control tend to be less lateralized in older adults than younger by functional neuroimaging study of cognitive performance (Cabeza 2002; Dolcos et al. 2002). The larger MI100 level in right-hemisphere for younger and comparable MI100 level for both hemisphere in older listeners also support the right hemi-aging model, which suggests that the right hemisphere shows greater age-related decline than the left hemisphere (Brown and Jaffe 1975). Age-related asymmetry reductions, i.e., increases in left-hemisphere processing, may reflect functional compensation. Dolcos et al. (2002) investigated aging effects on a letter-matching task with varying difficulty levels, and it suggested that older adults might benefit from bilateral processing at different task complexity levels. However, for younger adults, unilateral processing was sufficient enough in most cases. The present study extends the asymmetric reduction hypothesis to cortical processing of continuous speech for older listeners and suggests a bilateral compensation mechanism for older listeners in cocktail party listening conditions.

## CONCLUSION

Mutual information analysis provides a robust non-linear approach towards investigations of cortical representations of continuous speech. The mutual information representation has higher predictive power for behavioral measures compared to linear representations. Using this novel approach, the current results show that with aging, the cortical response to speech is not only larger in amplitude but also redundant in information. Finally, the late response component (∼200 ms latency) may be an important biomarker for older listeners, associated with both behavioral inhibition and speech comprehension.

## SOURCE DATA

Source data available at http://hdl.handle.net/1903/21184.

